# Agroforestry adoption and its influence on soil quality under smallholder maize production systems in western Kenya

**DOI:** 10.1101/2024.10.25.620197

**Authors:** H.T Nyuma, R Njoroge, A.N Otinga

## Abstract

Agroforestry, a sustainable land use practice was-introduced in western Kenya in the early 1990’s as a soil fertility replenishment strategy alongside other multiple benefits. Since then, effect of the practice on soil quality is seldom evidenced. Therefore, a study was conducted in the region to assess the effects of agroforestry adoption on soil quality under small holder maize production systems. A total of 120 soil samples were collected from two land use practices: agroforestry adoption (90) and non-agroforestry adoption (30) at 0-30 cm depth from two locations (Busia and Kakamega counties). On average, adoption of agroforestry significantly improved soil physicochemical properties compared to non-adoption of agroforestry. Bulk density (BD) reduced by 21% (from 1.4 to1.1g cm^−3)^ while SOC increased by 75% (0.8-1.4%), P by 80% (3.0-5.4 mg kg^−1^), exchangeable K^+^ by 256% (0.3-8.0 Cmolc kg^−1^), Ca^2+^ by 100% (1.0-2.0 Cmolc kg^−1^), S by 50%(0.2-0.3 mg kg^−1^), and Cu by 18% (2.8-3.3 mg kg^−1^).In reference to the soil environmental requirement for maize production, agroforestry adoption significantly increased K and Cu above the critical thresholds of 0.4 Cmolc kg^−1^ and 1.0 mg kg^−1^, respectively regardless of the study location or adoption practice. In addition, different agroforestry tree species had variable effect on soil properties. Sesbania and leucaena significantly influenced soil BD, clay, pH, Similarly, soil available P (4.3.-7.0 mg kg ^−1^), exchangeable K^+^ (0.4-0.7 cmolc kg^−1^), Mg (0.1-0.2 cmolc kg^−1)^, and Mn (13.5 – 25.2 mg kg^−1^) above non-agroforestry adoption at both locations, while calliandra significantly increased SOC in Kakamega only.

## 1.0 Introduction

Healthy soils are cardinal to sustainable agricultural production globally. Soil degradation as expressed by soil fertility decline presents numerous challenges to global food and nutrition security. Soil degradation affects 36–75 billion hectares of agricultural soils annually, hence threatening global food supply [1,2]. Soil degradation is linked to inappropriate agronomic practices, resulting in detrimental effects on soil ecosystem services including nutrient availability and carbon sequestration [3–5]. In Sub-Saharan Africa (SSA), soil degradation affects more than 40 million hectares of arable lands and the livelihood of more than 120 million farming households [4]. In Kenya, soil degradation affects more than 30% of the country’s arable land [6,7] with observed deficiencies of nitrogen (N), phosphorus (P) and zinc (Zn) and low organic carbon [8].

Maize is an important staple food crop for more than 75% of Kenya’s population, its production by smallholder agriculture systems is characterized by soil fertility decline, intensive cultivation, and low productivity [9]. Soil fertility loss in the country is triggered by an increasing human population, climate variability and unsustainable land use practices; showed a 30-60% decline in the yields of maize. The situation is further worsened by soil nutrient imbalances, resulting in the deficiency and or toxicity of essential macro and micronutrients [8,10]. This affects agricultural productivity, hence the nutritional insecurity of the country [11,12]. Practically, access to and the affordability of inorganic inputs, are among the constraints affecting agricultural productivity in most parts of the country[13].

Therefore, Alternative soil replenishment strategies are inevitable for sustainable agricultural productivity in the country. Agroforestry is a sustainable land use practice that contributes to sustainable food production, and the amelioration of environmental quality through the enhancement of soil ecosystem functions, [14–16]. The adoption of fast-growing agroforestry trees such as calliandra, gliricidia, leucaena, and sesbania commonly known as fertilizer trees are noted for their multiple functions such as the provision of fodder [17]; biofuel [18,19] soil conservation and carbon sequestration [20,21]. Studies by [22,23] highlighted the ecosystem benefits of these leguminous species as organic amendments, due to rapid decomposition and rates of nutrient release from their biomass. Gupta et al. [21] reported the amelioration of ecosystem functions and the restoration of degraded soils under agroforestry across different agroecological zones in tropical regions of the world. However, the adoption of agroforestry for soil fertility management specifically for improved yields of food crops in Kenya is low, usually attributed to land size, limited knowledge of the management of exotic agroforestry species, and ineffective extension services [24]. Reducing land size, and lack of training are reported as other challenges affecting the adoption of agroforestry among farming communities in SSA [18,25].

Reducing soil fertility gaps through agroforestry interventions has the potential to rejuvenate degraded agricultural land and improve household income and food security. Despite the adoption of agroforestry by smallholder farmers in western Kenya, benefits of the technology on soil health remain unclear. Therefore, the aim of the study was to evaluate the effects of agroforestry adoption on soil quality under smallholder maize production systems in western Kenya. It was hypothesized that agroforestry adoption enhances soil quality for optimum maize production.

## 2.0 Methods

### 2.1 Study area and site selection

The study was conducted from January – July 2023 in Butsotso ward, Lurambi sub county of Kakamega County, located at longitude 0°17’3.19”N and latitude 34°45’8.24”E and in Amukura, Chakol North ward, Teso South sub county of Busia County located at longitude 0°25’59.99”N, latitude 34°08’60.00” E). Both study locations are situated in the Midland (LM) Zone 2-3 and Upper Midland (UM) Zone 4 Agro-ecological zones in western Kenya[26] Figure 1.

**Figure 1:**
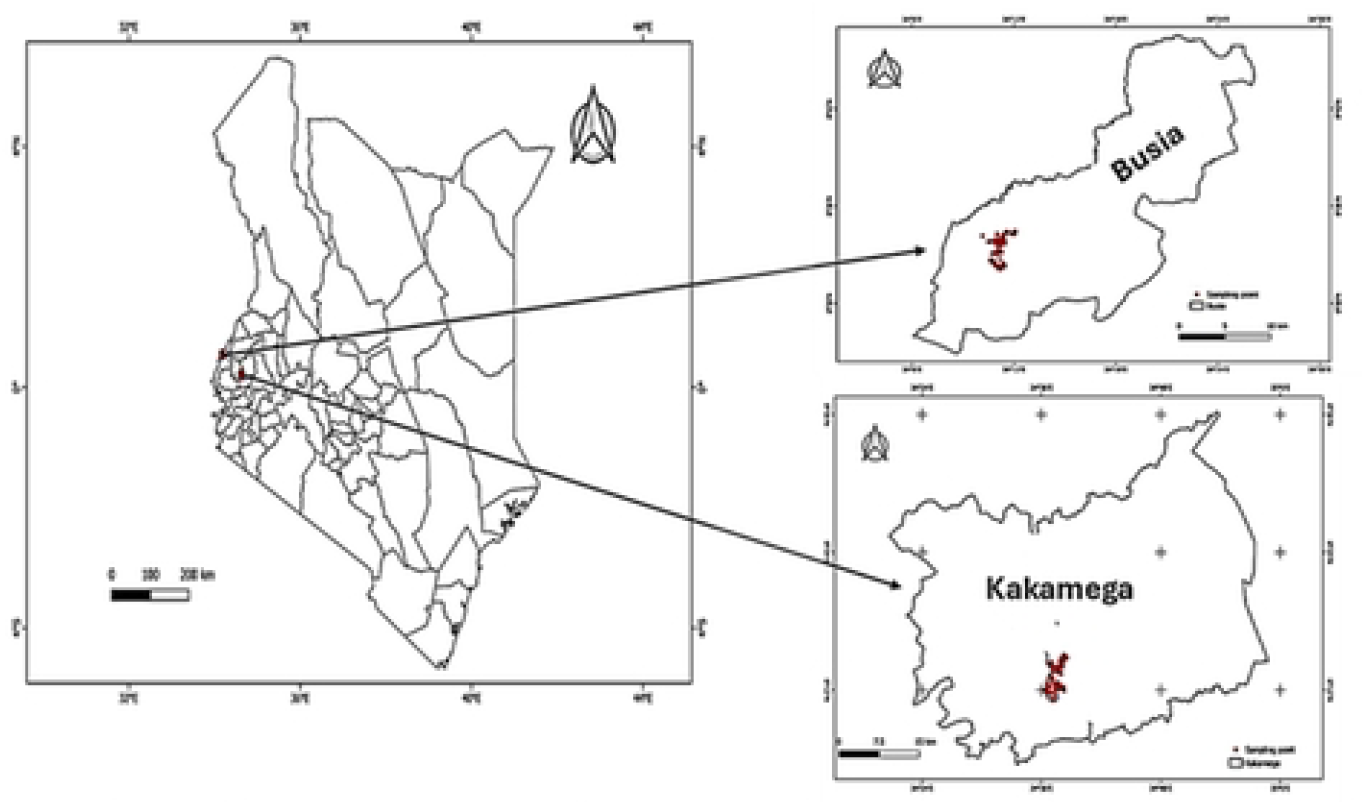
A map of the study locations and sampling points at the study locations in western Kenya.

Rainfall in western Kenya is above average, showing a bimodal distribution with two distinct wet seasons. The long rains occur from March-May and short rains from October-December. Average rainfall during long rains is 1000 - 1200 mm while short rains is 500 – 800 mm annually (27). Soils in the study area are highly weathered and acidic in nature with low inherent soil fertility status (low N and P, as well (Ca^2+,^ Mg^2+^ and K^+^. The soils are broadly classified as Ferralsols and Nitisols in Busia and Kakamega, respectively [28,29].

### 2.1 Study design

The study sites were purposely selected based on similarity in agro-climatic conditions and the dominance of agroforestry activities [30]. The study involved a field survey using an open-ended questionnaire per household and collection of soil samples for laboratory analyses to assess the effects of adoption practices (agroforestry adoption and non-adoption), and selected agroforestry tree species Calliandra (*Calliandra calothyrsus*) Leucaena (*Leucaena leucocephala*) and Sesbania (*Sesbania sesban (L*.*) Merr*.) on soil quality for maize production.

### 2.2 Data collection

#### 2.2.1 Soil sampling

A total of 120 soil samples were collected from 60 locations in Busia and Kakamega counties. To make a comparison between changes in soil characteristics under agroforestry and non-agroforestry practices, given that study sites are predominantly under agroforestry practices, 75% (90) of samples were collected from agroforestry adopters while 25% of the samples (30) were collected from non-agroforestry adopters, following sampling guidelines prescribed by [31].

Soil samples under agroforestry adoption were randomly collected within 1m radius from the base of agroforestry trees in three directions and properly mixed to form composite samples. Sampling in non-agroforestry adoption plots was done randomly from farm fields, however, in fields where crops were established, soil samples were collected between crop rows to ensure minimum interference with their roots. All samples were properly labelled, packaged in khaki paper bags and transported to the Department of Soil Science Laboratory at the University of Eldoret (111-115 km) away from the study sites, for physicochemical analyses.

#### 2.3 Field and laboratory soil measurements

All field sampling and laboratory analytical procedures were conducted in accordance with [31] soil manual. Particle size distribution was determined by the Bouyoucos hydrometer [32]. Soil bulk density (BD) was determined insitu by the core ring method [33]. Soil pH was determined in 1:2.5 H_2_O suspension using a glass electrode pH meter model: HI 2211, Hanna instruments). Total N was determined by Kjeldahl method soil organic carbon (SOC) was determined by potassium dichromate method. Available P was determined following the Olsen method, Ca (cmol_c_ kg^−1^), and Mg (cmolc kg^−1^) were analyzed by atomic absorption spectroscopy, K (cmolc kg^−1^) was analyzed by flame photometry and micronutrients: Cu (ppm), Fe (ppm), Mn (cmolc kg^−1^) and Zn (ppm) were determined by atomic absorption spectroscopy after ethylenediaminetetraacetic acid (EDTA) extractions as described in Table 1.

**Table 1:**
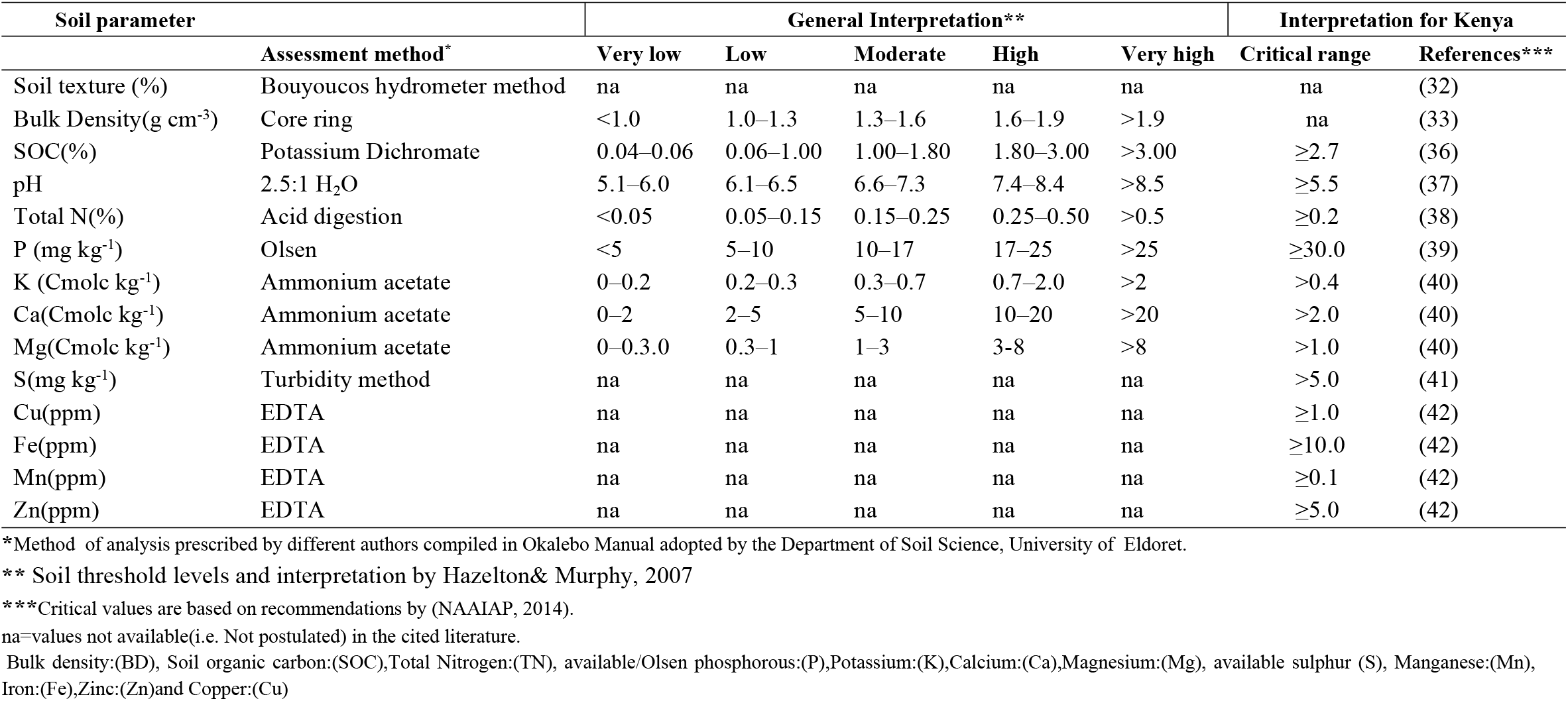
Soil analysis procedures and threshold levels postulated by [34] versus critical values for maize growth according to the Kenya national soil report [35].

#### 2.4 Data analysis

Data was organized in Microsoft EXCEL, 2013 version and subjected to statistical analysis using R version 4.2.2 (R Core Team, 2023). Significant differences (p < 0.05) in soil properties were assessed using linear mixed-effects models (lme4 package). Least significant difference (LSD) pairwise post hoc tests were performed for means comparison at p < 0.05, where factors were identified as statistically significant for adoption practices and study locations. A one-way ANOVA was conducted to determine the effects of agroforestry tree species on soil characteristics, and means were declared significant at (p < 0.05) according to Tukey’s Honesty Test. Pearson correlation coefficients were calculated to assess the relationship between soil mineral elements and physicochemical properties. Soil quality indicators (BD, SOC pH, exchangeable bases, macro and micronutrients) were assessed as prescribed by [31]. Thereafter, the results of soil physicochemical characteristics were interpreted using thresholds previously compiled and summarized by [34] and compared against the critical nutrient levels for maize growth in Kenya recommended by [35].

## 3.0 Results

### 3.1 Influence of agroforestry adoption and study location on soil characteristics

Findings from the study showed significant difference in soil physicochemical properties between adoption practices and study locations Table 2. Generally, agroforestry adoption significantly influenced BD(1.3-1.2 g cm^−3^), SOC(0.8-1.4%), P(3.0-5.4 mg kg^−1^),K(0.c-0.8 |Cmolc kg^−1^)Ca(1.0 -2.0 Cmolc kg^−1^),S(0.2-0.3 mg kg^−1^)Cu(2.8-3.3 mg kg^−1^), and Mn(18.3-25.2 mg kg^−1^) over non-agroforestry adoption. Between the study locations, the results showed that soils in Kakamega had lower soil BD (1.3 g cm^−3^), higher clay (19.2%), SOC (1.5%), TN (0.2%),K(0.8 Cmolc kg^−1^),Ca(2.2 Cmolc kg^−1^),Mg(0.4 Cmolc kg^−1^), Cu(3.7 mg kg^−1^), Fe(3.7 mg kg^−1^),and Mn(27.1 mg kg^−1^) than soils in Busia Table 2. Mean BD in Busia (1.3 g cm^−3^) was significantly (P≤ 0.05) higher than that of Kakamega (1.2 g cm^−3^). Similarly, agroforestry adoption significantly reduced BD at both locations, with an observed decrease from 1.4 to 1.2 g cm^−3^, indicative of an 11% reduction in BD Table 2. Soil clay content at both study locations ranged from 2 to 40% with a mean of 14% but did not differ between adoption practices at both locations. However, clay content in soils at both study locations differed significantly (P≤ 0.05). Soils from Kakamega recorded a mean clay content of 19%, which was 43% higher than clay content in soils from Busia County Table 2. Soil organic carbon differed significantly (P≤0.05) between the study locations and between adoption practices (Table 2). Soil organic carbon ranged from 0.1 and 3.6% with a mean of (1.0%) in Busia and (1.5%) in Kakamega. Significant (P≤0.05) increases in SOC under agroforestry adoption 1.0-1.1% in Busia and 0.9-1.7% in Kakamega as observed represented 5% and 49% increase in SOC in Busia and Kakamega, respectively Table 2.Soil pH recorded in the study ranged from 4.5-6.9, and in the range of strongly acidic (4.5-5.5) to slightly acidic (5.6-6.5) according to (35). Differences in soil pH was observed at both locations and adoption practices Table 2. Farms adopting AF showed significant reduction in pH from (6.3-5.5) at both locations Table 2.Within the study locations, it was revealed that agroforestry adoption significantly reduced BD(1.3-1.2 g cm-3), and TN(0.2-0.3%),but increased SOC(0.7-1.1%),K(0.4-0.5 Cmolc kg^−1^),Ca(0.8-1.0 Cmolc kg^−1^),Cu(2.3-2.7 ppm), and Mn(14.0-22.0 ppm). However, pH, TN, and Zn decreased significantly under agroforestry adoption in Busia, but clay content, P, Mg, S and Fe did not differ statistically between adoption practices in Busia. Similar patterns were observed in Kakamega, where agroforestry adoption significantly decreased BD(1.3-1.2g cm^−3^), and TN(0.2-0.4%),but increased SOC(1.0-2.0%)K(1.0-0.3 Cmolc kg^−1^),Ca(1.1-3.0 Cmolc kg^−1^),Cu(3.3-3.9 ppm),and Mn(23.0-29.0 ppm), while pH, Mg, S, Fe and Zn did not differ between adoption practices in Kakamega Table 2.

**Table 2:** Mean physico-chemical characteristics of soils as influenced by agroforestry adoption and study location.

| Soil properties | Busia |  |  | Kakamega |  |  |
| --- | --- | --- | --- | --- | --- | --- |
|  | Adopters | Non adopters | Mean Busia | Adopters | Non adopters | Mean Kakamega |
| BD (g cm<sup>-3</sup>) | 1.2(±0.1)b | 1.3(±0.1)a | 1.3(±0.1) | 1.1(±0.1)a | 1.3(±0.1)b | 1.2(±0.1) |
| Clay (%) | 8.5(±1.0)a | 7.3(±1.0)a | 8.2(±0.1) | 20.4(±1.1)a | 16.0(±2.0)b | 19.2(±0.1) |
| SOC (%) | 1.1(±0.1)a | 0.7(±0.1)a | 1.0(±0.1) | 2.0(±0.1)a | 1.0(±0.2)b | 1.5(±0.1) |
| pH (H<sub>2</sub>O) | 5.8(±0.1)b | 6.3(±0.1)a | 5.8(±0.1) | 5.5(±0.1)a | 5.7(±0.1)a | 5.5(±0.1) |
| TN (%) | 0.2(±0.1)b | 0.3(±0.1)a | 0.1(±0.1) | 0.2(±0.1)b | 0.4(±0.1)a | 0.2(±0.1) |
| P (mg kg<sup>-1</sup>) | 5.5(±0.3)a | 4.3(±0.5)a | 5.2(±0.1) | 5.3(±0.2)a | 1.6(±0.5)b | 4.3(±0.3) |
| K (Cmolc kg<sup>-1</sup>) | 0.5(±0.1)a | 0.4(±0.1)b | 0.5(±0.1) | 1.0(±0.1)a | 0.3(±0.1)b | 0.8(±0.1) |
| Ca (Cmolc kg<sup>-1</sup>) | 1.0(±0.1)a | 0.8(±0.1)b | 1.0(±0.1) | 3.0(±0.2)a | 1.1(±0.3)b | 2.2(±0.1) |
| Mg (Cmolc kg<sup>-1</sup>) | 0.2(±0.1)a | 0.1(±0.1)a | 0.1(±0.1) | 0.4(±0.1)a | 0.3(±0.1)a | 0.4(±0.1) |
| S(mg kg<sup>-1</sup>) | 0.2(±0.1)a | 0.2(±0.1)a | 0.3(±0.1) | 0.3(±0.1)a | 0.2(±0.1)a | 0.2(±0.1) |
| Cu(ppm) | 2.7(±0.1)a | 2.3(±0.1)b | 2.6(±0.1) | 3.9(±0.1)a | 3.3(±0.1)b | 3.7(±0.1) |
| Fe(ppm) | 2.5(±0.1)a | 2.4(±0.1)a | 2.5(±0.1) | 3.7(±0.1)a | 3.8(±0.3)a | 3.7(±0.1) |
| Mn(ppm) | 22.0(±0.5)a | 14.0(±1.0)b | 20.0(±1.0) | 29.0(±0.8)a | 23.1(±1.2)b | 27.1(±1.0) |
| Zn(ppm) | 0.9(±0.1)b | 1.2(±0.1)a | 1.0(±0.1) | 1.6(±0.2)a | 1.1(±0.3)a | 1.4(±0.1) |

Means with different letters within a row indicate significant differences ( $p \leq 0.05$ ) for columns representing different factors (i.e. agroforestry adoption practice, and adoption \* location).

Total N-content ranged from 0.2 – 0.3% between study locations and from 0.2-0.4% adoption practices. Results of the study revealed that agroforestry adoption significantly influenced TN content in Busia (1.0 -1.1%). On the contrary, TN decreased with agroforestry adoption from 0.4-0.2% in Kakamega county Table 2. Soil available P did not differ between the study locations, but were statistically different among adoption practices in Kakamega, and did not differ among adoption practices in Busia Table 2. Soil available S in the study did not differ between the study location but varied significantly among adoption practices in both counties. Changes in S as affected by agroforestry adoption was observed to increase from 0.2 -0.3 mg kg^−1^ as shown in Table 2.Soil exchangeable Ca^2+^ in the study ranged from 0.4 to 6.1 cmolc kg^−1^and differed significantly (P≤0.05) between the study locations and between adoption practices Table 2. Exchangeable K^+^ in soils from both locations ranged from (0.2 – 2.0 cmol_c_ kg^−1^) and was rated as low (< 0.2 cmol_c_ kg^−1^) to medium (2.0 -3.5 cmol_c_ kg^−1^) according to (35). Exchangeable K^+^ differed between the study locations with soils from Busia recording the highest exchangeable K^+^ content (0.8 cmol_c_ kg^−1^). It was further revealed that agroforestry adoption significantly augmented soil exchangeable K^+^ from 0.3 - 1.0 cmol_c_ kg^−1^ over non-agroforestry adoption at both locations Table 2. Exchangeable Mg^2+^ in the study ranged from (0.1 to 2.0 cmol_c_ kg^−1^). It was observed that Mg^2+^ contents differed significantly (P≤ 0.05) between the study location and adoption practices Table 2. Micronutrient contents of soils in the study differed significantly (P<0.05) between the study locations and between adoption practices Table 2. Agroforestry increased Cu (2.4-4.0 mg kg^−1^), Mn (14.0 – 29 mg kg^−1^) and Zn (0.9 -1.6 mg kg^−1^) over the control in the study. Whereas Fe did not differ between the study locations and adoption practices Table 2.

### 3.2 Spatial distribution soil properties as influenced by agroforestry adoption in Busia and Kakamega

#### 3.2.1 Distribution of SOC and pH in soils under smallholder maize production

Average soil pH in Busia (5.8) and Kakamega (5.5) were within the critical range (5.5) for maize production. However, soils under non-agroforestry adoption recorded a higher pH than those under agroforestry adoption with more than 50% of all soil samples from both locations recorded a pH of > 5.5 Figure 2. The distribution of SOC in the study revealed that about 88% of all samples (both adopters and non-adopters) recording low SOC contents (< 2.0%), below the critical threshold for maize production in Kenya Figure 2.

**Figure 2:**
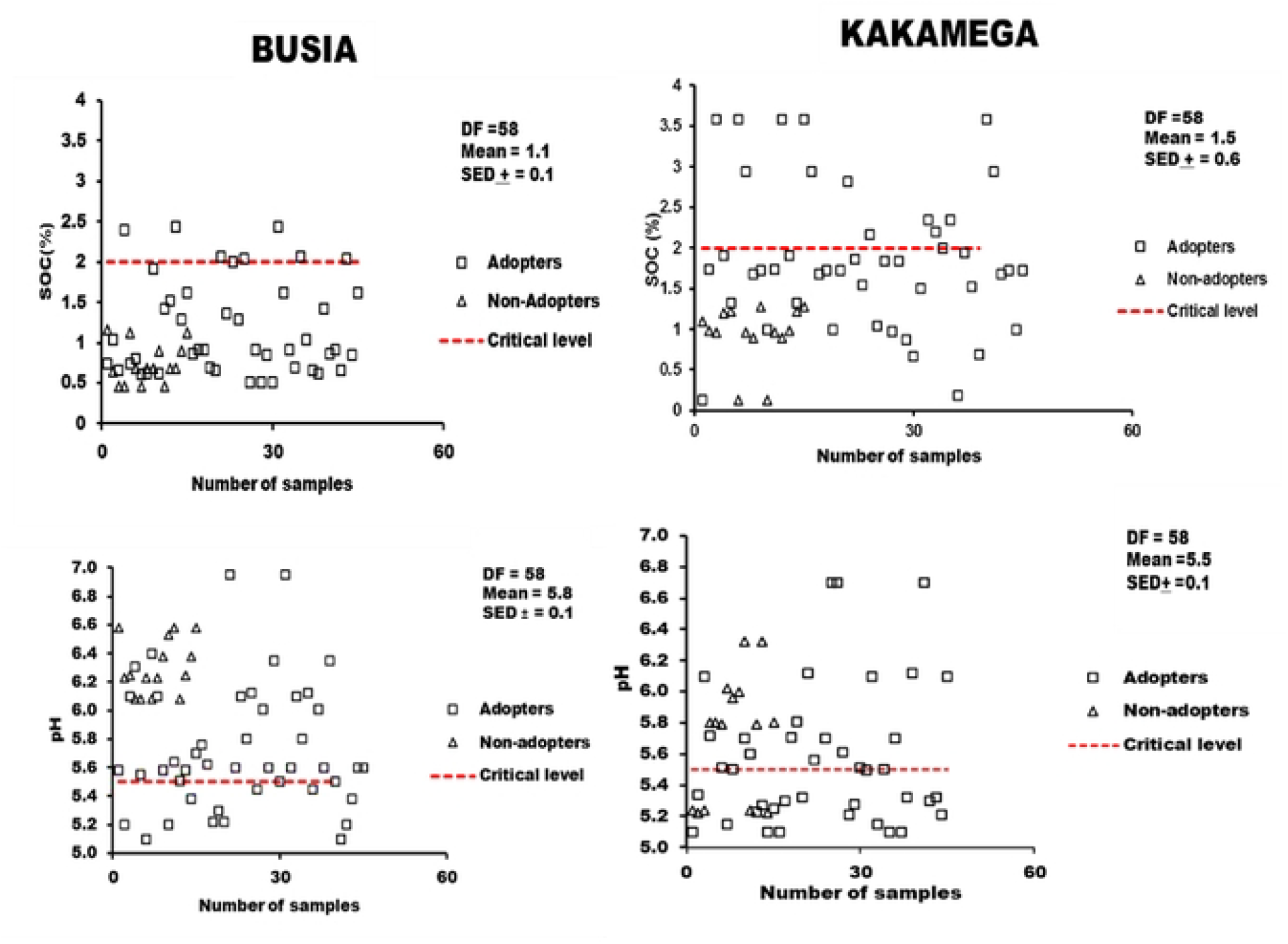
Spatial distribution of SOC and pH and their respective critical levels in Busia and Kakamega.

#### 3.2.2 Distribution of macronutrients in soils under small holder maize production systems in Busia and Kakamega

Distribution of soil -N showed that more than 80% of soil samples from both locations were below the critical threshold for maize production Figure 3. Soil available P at both locations were critically low. All samples under agroforestry and non-agroforestry adoption in the study were recorded below the critical P level (<10.0 mg kg^−1^) for maize production Figure 3. Mean available S in the study (0.3 mg kg^−1^) was below the critical threshold (0.5 mg kg^−1^) for maize Figure 3. Available S did not different between the study locations, however all samples at both locations were recorded below the critical threshold Figure 3.

**Figure 3:**
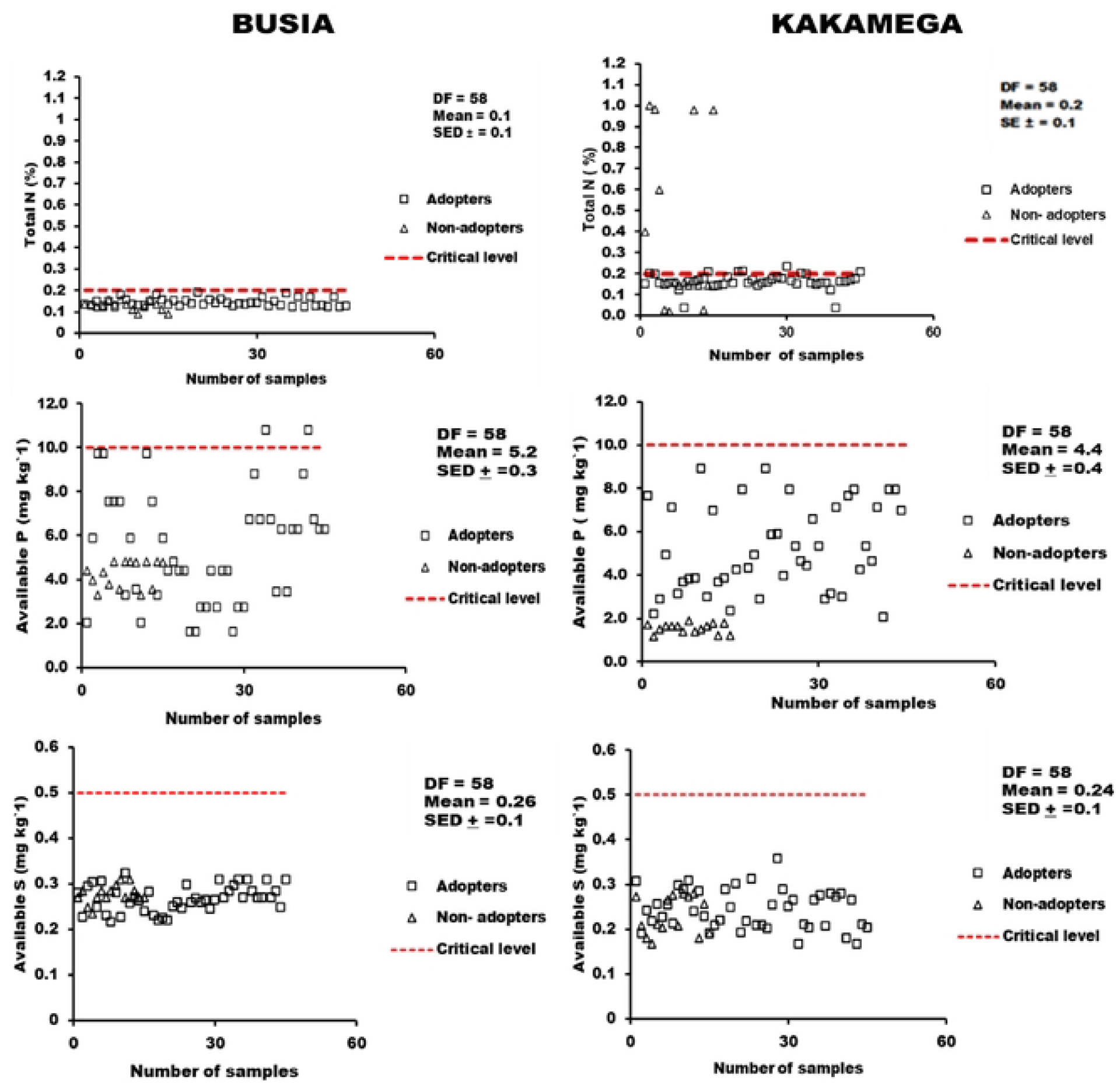
Distribution and availability of macronutrients among agroforestry adopters and non-adopters in Busia and Kakamega counties.

#### 3.2.3 Distribution of exchangeable bases in soils under small holder maize production systems in Busia and Kakamega

It was observed that more than 70% and of the soil samples from Kakamega and 93% from Busia were below the critical nutrient level for Ca (2.0 cmol_c_ kg^−1^) for maize production Figure 4. The results also showed that about 75% of soil samples from both locations were above the critical level for K^+^ (0.4 cmol_c_ kg^−1^) and differed significantly between adoption practices. Increase in exchangeable Mg^2+^ at both locations was influenced by agroforestry, however all samples from both sites were below the critical threshold (1.0 cmol_c_ kg^−1^) for maize production Figure 4.

**Figure 4:**
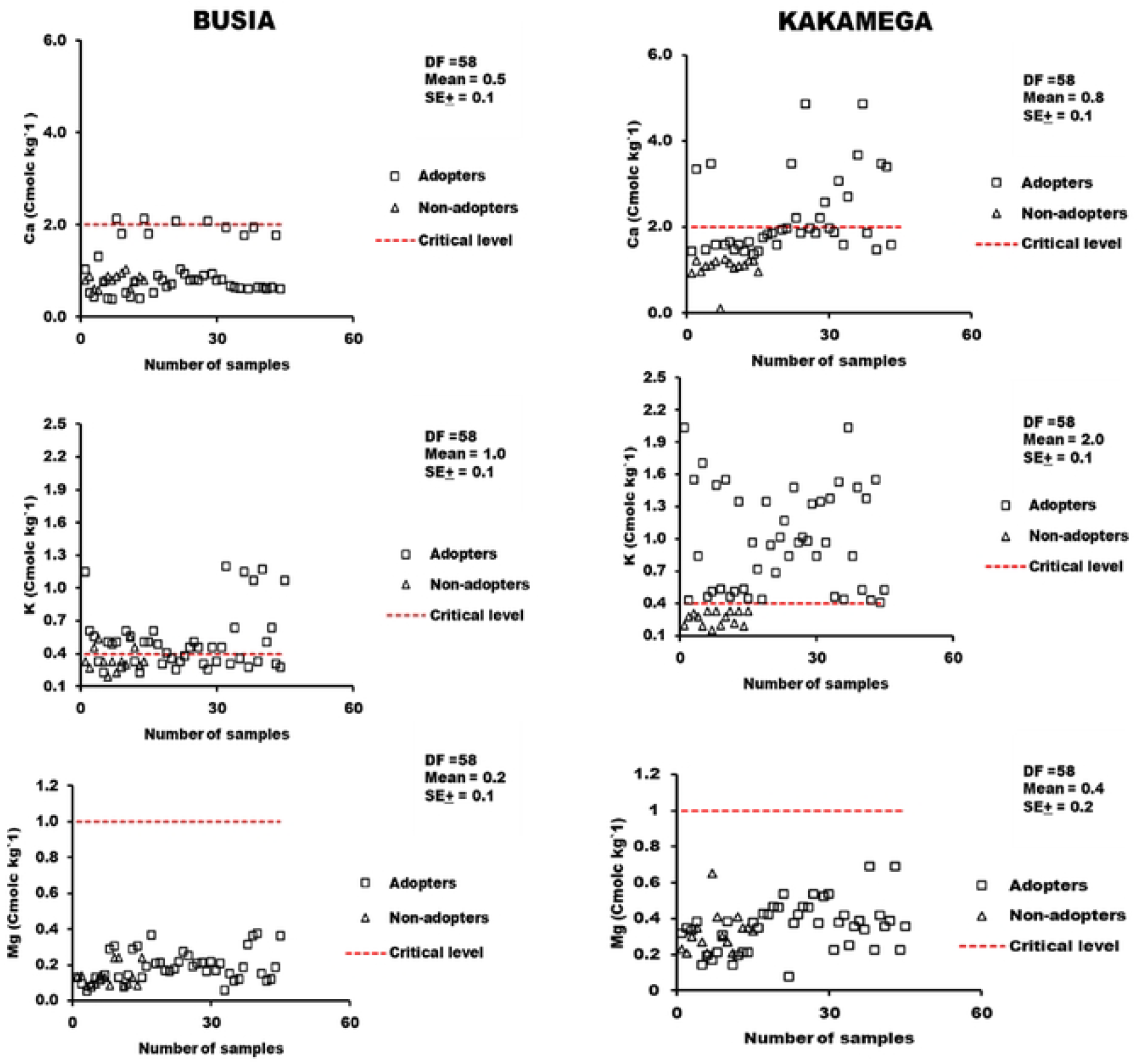
Distribution and availability of soil exchangeable bases in soils under small holder maize production in Busia and Kakamega

##### Distribution of micronutrients in soils under small holder maize production systems in Busia and Kakamega

Availability of micronutrients in both sites and under adoption practices were above the critical level (1.0 mg kg^−1^) for maize production, Figure 5. On the contrary, all samples from Busia recorded low levels of Fe, Mn and Zn contents below their respective critical levels for maize production, while only 40% of the samples from Kakamega were above the critical threshold for Mn Figure 5.

**Figure 5:**
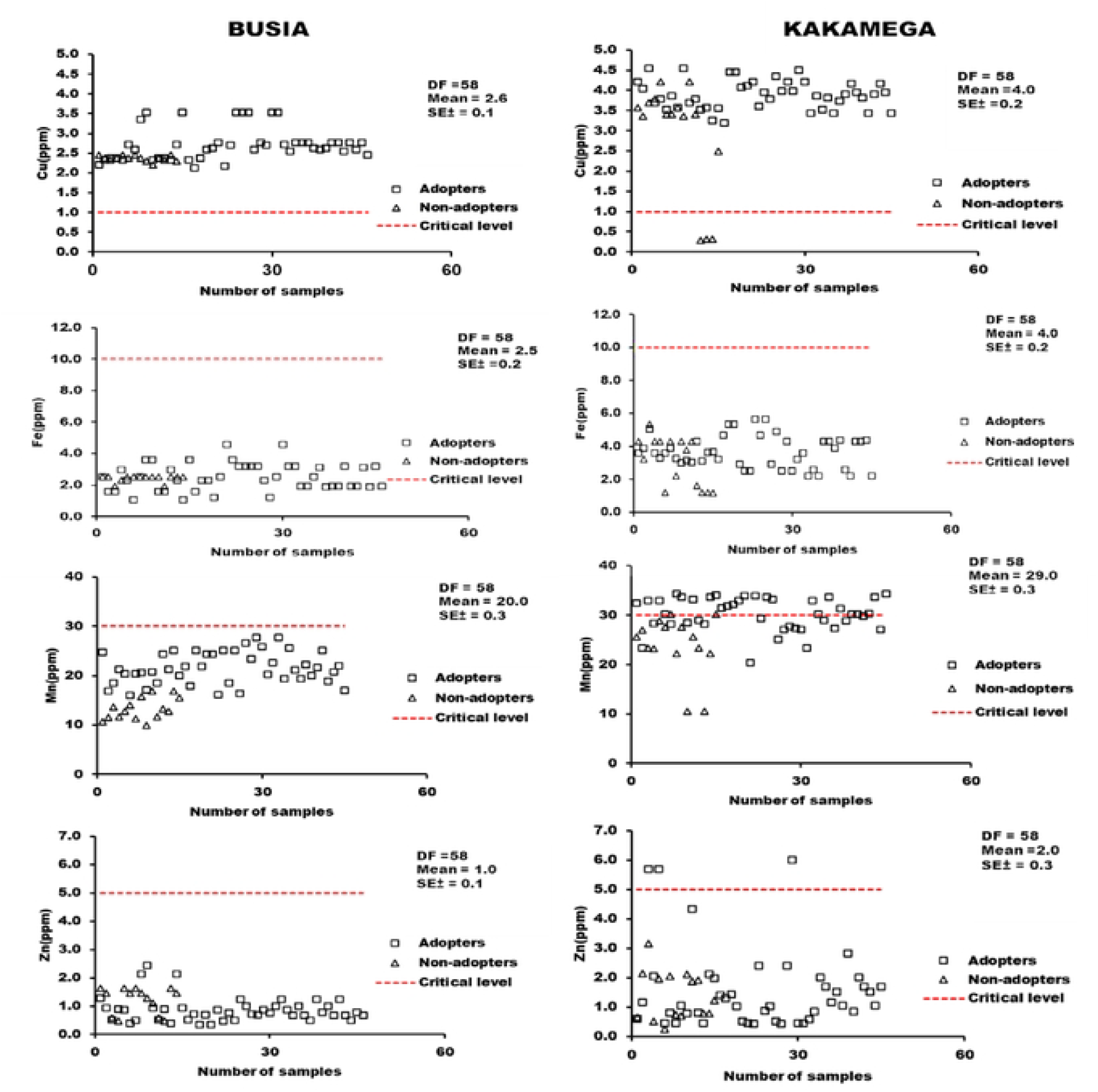
Distribution of available micronutrients in soils under agroforestry and non-agroforestry adoption in Busia and Kakamega

### 3.7 Effects of selected agroforestry trees species on soil chemical properties

Agroforestry trees species had variable effects on the physico- chemical characteristics of soils at the study locations with leucaena and sesbania showing the potential to enhance macronutrients and micronutrients Table 3. Generally, agroforestry tree had positive influence on the overall characteristics of soils as observed in. Significant increase in soil available P (4.3.-7.0 mg kg ^−1^), exchangeable K^+^ (0.4-0.7 cmolc kg^−1^), Mg (0.1-0.2 cmolc kg^−1)^, and Mn (13.5 – 25.2 mg kg^−1^) were recorded in soils under leucaena and sesbania above non-agroforestry adoption in Busia. Similar patterns were observed in Kakamega; however, it was observed that calliandra increased SOC from 0.9 – 2.1% over non-agroforestry in Kakamega Table 3. Results from the study showed that sesbania significantly influenced soil BD, clay, pH, available P, and exchangeable K^+^, contents of soils in Busia above the other tree species.

**Table 3:**
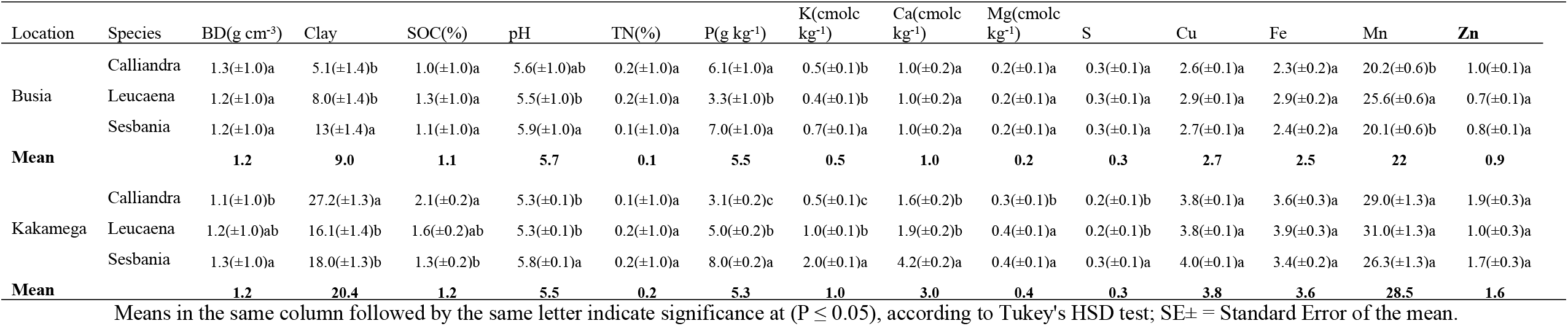
Effects of agroforestry tree species on the chemical properties of soils in Busia and Kakamega counties of western Kenya.

| Location | Species | BD(g cm<sup>-3</sup>) | Clay | SOC(%) | pH | TN(%) | P(g kg<sup>-1</sup>) | K(cmolc kg<sup>-1</sup>) | Ca(cmolc kg<sup>-1</sup>) | Mg(cmolc kg<sup>-1</sup>) | S | Cu | Fe | Mn | Zn |
| --- | --- | --- | --- | --- | --- | --- | --- | --- | --- | --- | --- | --- | --- | --- | --- |
| Busia | Calliandra | 1.3(±1.0)a | 5.1(±1.4)b | 1.0(±1.0)a | 5.6(±1.0)ab | 0.2(±1.0)a | 6.1(±1.0)a | 0.5(±0.1)b | 1.0(±0.2)a | 0.2(±0.1)a | 0.3(±0.1)a | 2.6(±0.1)a | 2.3(±0.2)a | 20.2(±0.6)b | 1.0(±0.1)a |
|  | Leucaena | 1.2(±1.0)a | 8.0(±1.4)b | 1.3(±1.0)a | 5.5(±1.0)b | 0.2(±1.0)a | 3.3(±1.0)b | 0.4(±0.1)b | 1.0(±0.2)a | 0.2(±0.1)a | 0.3(±0.1)a | 2.9(±0.1)a | 2.9(±0.2)a | 25.6(±0.6)a | 0.7(±0.1)a |
|  | Sesbania | 1.2(±1.0)a | 13(±1.4)a | 1.1(±1.0)a | 5.9(±1.0)a | 0.1(±1.0)a | 7.0(±1.0)a | 0.7(±0.1)a | 1.0(±0.2)a | 0.2(±0.1)a | 0.3(±0.1)a | 2.7(±0.1)a | 2.4(±0.2)a | 20.1(±0.6)b | 0.8(±0.1)a |
| **Mean** |  | **1.2** | **9.0** | **1.1** | **5.7** | **0.1** | **5.5** | **0.5** | **1.0** | **0.2** | **0.3** | **2.7** | **2.5** | **22** | **0.9** |
| Kakamega | Calliandra | 1.1(±1.0)b | 27.2(±1.3)a | 2.1(±0.2)a | 5.3(±0.1)b | 0.1(±1.0)a | 3.1(±0.2)c | 0.5(±0.1)c | 1.6(±0.2)b | 0.3(±0.1)b | 0.2(±0.1)b | 3.8(±0.1)a | 3.6(±0.3)a | 29.0(±1.3)a | 1.9(±0.3)a |
|  | Leucaena | 1.2(±1.0)ab | 16.1(±1.4)b | 1.6(±0.2)ab | 5.3(±0.1)b | 0.2(±1.0)a | 5.0(±0.2)b | 1.0(±0.1)b | 1.9(±0.2)b | 0.4(±0.1)a | 0.2(±0.1)b | 3.8(±0.1)a | 3.9(±0.3)a | 31.0(±1.3)a | 1.0(±0.3)a |
|  | Sesbania | 1.3(±1.0)a | 18.0(±1.3)b | 1.3(±0.2)b | 5.8(±0.1)a | 0.2(±1.0)a | 8.0(±0.2)a | 2.0(±0.1)a | 4.2(±0.2)a | 0.4(±0.1)a | 0.3(±0.1)a | 4.0(±0.1)a | 3.4(±0.2)a | 26.3(±1.3)a | 1.7(±0.3)a |
| **Mean** |  | **1.2** | **20.4** | **1.2** | **5.5** | **0.2** | **5.3** | **1.0** | **3.0** | **0.4** | **0.3** | **3.8** | **3.6** | **28.5** | **1.6** |

Means in the same column followed by the same letter indicate significance at ( $P \leq 0.05$ ), according to Tukey's HSD test; SE± = Standard Error of the mean.

On the overall, it was observed that pH, and K increased above their respective critical values under the canopy of sesbania., while leucaena increased Mn of soils in Busia Table 3. In Kakamega, clay, BD, and SOC were significantly increased under the canopy of calliandra, whereas sesbania increased available P, S, pH, exchangeable Ca^2+^, and K^+^. It was also observed that pH, K^+^, and Ca^2+^ in soils under the canopy of sesbania increased above the critical values for maize production Table 3

## 4.0 Discussion

### Influence of agroforestry adoption soil characteristics

Changes in soil characteristics as observed in the study could be ascribed to differences in soil biogeochemical characteristics between the study sites and land use practices. Lower soil bulk density of (1.2 g cm^−3^) under agroforestry systems can be attributed to organic matter build up as the result of litter fall and high turnover of finer roots [43]. Long term accumulation of organic matter through litter fall has the potential to buffer soil against the impact of raindrop, compaction and wind erosion, hence a lower bulk density[44]. These findings agree with [45]who reported that agroforestry trees reduced soil BD from (1.9-1.3 g cm^−3^) within 1 meter distance under the canopy of agroforestry trees in Ethiopia, Similarly, [46,47] reported improvement in soil bulk density and overall physical fertility of soils under agroforestry systems. Findings by [46,48] suggested agroforestry as an effective strategy for soil conservation due to its ability to reduce soil erosion and enhance soil physicochemical characteristics. Results of the current study is contrary to the findings of [49]who reported no difference in soil BD between agroforestry adopters and non-adopters.

The findings of [44] concord with findings by [47,50–52] who reported significant increase in soil bulk density associated with agroforestry practices, probably due to increased compaction of soil with machineries compared to traditional tillage practices. However, [53,54] observed that addition of organic matters from decomposing leaves and wood debris are important to improving soil physical properties, including BD.

Findings of this study suggests that agroforestry did not significantly influence soil clay content, although the clay content under agroforestry adoption was higher than those under non-agroforestry adoption Table 2. However, significant differences in clay content between the study sites can be attributed to the inherent nature of the soil physical properties, land clearing, low return of organic matter to the soil, and tillage practices were observed Table 2. Results of the current study corroborate the findings of [55] who reported no significant difference in clay content under agroforestry in the central highlands of Kenya.

Soil organic carbon content in the study (0.9-1.7%) was below the critical threshold (> 2.0%) for maize production in Kenya and emphasizes the need for increasing organic carbon stock at both locations. Low SOC status in the study can be is caused by low return of biomass given competing priorities for biomass such as fodder and fuel at household level [56]Other researchers [57,58] mentioned education, biophysical limitations, agronomic practices, and limited extension services as causes of low level of awareness on SOC loss under small holder agricultural systems. However,[59] reported increased SOC under gliricidia-maize intercropping systems over monoculture maize due to the decomposition of organic matter, hence the need for incorporating organic matter in soils for sustainable production.

Contrary to the findings of the current study Suleiman [60] reported no change in SOC contents in soils under agroforestry influence of agroforestry due to extreme drought, which affected litterfall and biomass accumulation.

Soil pH in the study decreased in fields adopting agroforestry due to the decomposition of organic matter and the release of organic acids. Findings of this is like findings by [61] who reported 40% decrease in soil acidification under agroforestry. Soil acidity (< 5.0) reduces the availability of essential nutrient elements such as P, Ca and Mo become limiting due to the solubility of Al^+3^ and Fe^3^, especially under humid conditions [62,63]. Such conditions are the main factors affecting cereal yields, thus amelioration through liming and appropriate management practices were recommended by [62,63].

Significantly decrease of TN contents in soils under agroforestry adoption at both study locations can be linked to decomposition of organic matter, and the leaching of soil N due to high rain fall in the western region of Kenya[64,65]. However, few cases of the use of calcium ammonium nitrate (CAN) as reported by farmers could be responsible for increase in TN among non-adopters. Findings from the study are contrary to findings by [66,67] who observed 9–19% increase in soil N content as well as the positive effects of agroforestry adoption on soil chemical fertility from (0-30 cm depth) under soybean agroforestry-system. Riyadh et al. ([68] recommended agroforestry as an alternative and cheaper source of crop nutrients to sustain smallholders’ productivity amidst the increasing costs of fertilizer inputs.

Available P in the study was limiting element with more than 50% of the samples from both sites below the critical P range (30 mg kg^−1^). Like in most parts of SSA, phosphorus deficiency is caused by immobilization and fixation in the presence of Fe and Al^3+^ under acidic conditions [67]. However, an overall 45% increase in soil available P due to agroforestry in this study is an indication that agroforestry adoption has the potential to enhance soil available P requires further investigation [69]. Findings of the study confirms reports by [70] who observed an 11% increase in soil P under agroforestry practices compared with monocropping systems.

Available S in the study was below the critical threshold for maize production, and can be attributed to soil type, organic carbon content, climatic, clay and N contents [71]. These findings concord with reports by [72] who suggested the need for including S into soil fertility management programs for small holder farming systems in SSA where fertilizer inputs mainly focus on the primary macronutrients (N,P,and K).

Low exchangeable Ca^2+^ in the study could be linked to management practices and clay mineralogy of soil in the study area [62,73].

[62,73] reported soil acidity and poor soil management practices as a leading cause of Ca^2+^ deficiency in western Kenya, hence the need for adopting sustainable management practices, such as the use of organic amendments. Findings of the study concord with findings reported by [74] who reported slight changes in exchangeable Ca^2+^concentrations between agroforestry and other land use systems. Contrary to findings of the study, [75] reported a 17-57% increase in exchangeable bases under agroforestry compared with treeless fields. Higher concentration of Ca^2+^, especially in the non-agroforestry plots, can be attributed to the use of inorganic fertilizer and agricultural lime, whose use is determined by the financial capability of few farmers, thus the need for a cheaper and sustainable source of Ca^2+^. [76] suggested agroforestry practices such as improved fallow as a sustainable mean to improve soil chemical fertility and the availability of basic cations.

Agroforestry adoption increased exchangeable K^+^ above the critical threshold which is in conformity with the findings of [8] who reported optimum concentrations of K^+^ in soils of western Kenya for maize. Potassium contents could be influenced the chemistry and clay mineralogy of soils. The enhancement of soil exchangeable K^+^ in this study is an important component of regenerative agriculture and has been recommended by several researchers [77– 79].Increased exchangeable K^+^ in degraded acid soils under agroforestry is evident of nutrient recycling in agro ecosystems have after the application of post-harvest cereal residues [66,74]. The findings of the current study concord with the findings of [66,74] who reported enhanced K^+^ availability in acid soils (Acrisols, Nitisols, and Ferralsols), which have notably poor K^+^-selective binding capacity and are low in potassium.

Exchangeable Mg^2+^ in all samples was rated as low (< 1.0 cmolc kg^−1^) for maize production, according to [35]. Findings from the study are contrary to the findings of [80] who reported a 33% decrease in exchangeable Mg^2+^ under agroforestry fallow system compared with other land use practices. Magnesium deficiency can be attributed to low organic matter content, soil acidity and crop removal of basic cations [63] with negative consequence on the quantity and quality of food production as Mg^2+^ has control over the photosystems of crop leaves [81].

The effects of agroforestry on Cu and Mn contents in soils as reported in the current study concord with the findings of [66,68]who reported increased soil micronutrients under agroforestry over the control plots, but Zn and Fe were the most deficient micronutrients in the study. The deficiency of Fe and Zn can be attributed to low SOC content and poor agronomic practices as observed in the current study. [82,83] observed that precipitation and increased adsorption to reactive surfaces such as soil organic matter and metal (hydr)oxides affect the availability of Zn due to the mineralization of organic matter or the formation of soluble organic Zn complexes [84]. (85) identified Zn as the most deficient micronutrient in arable soils across sub–Saharan Africa, affecting food and nutrition security, however they recommended agronomic biofortification micronutrient as a potential solution to ameliorate micronutrient deficiencies and improve crop productivity and nutritional quality of agricultural produce. Kaur et al. [78] reported (24.8% and 50.8%) increase in the total contents of Cu and Mn due to agroforestry, highlighting the role of agroforestry in nutrient cycling. These findings are in line with findings with works reported by [8] who linked high levels of micronutrient deficiency with malnutrition among children in Kenya, hence highlighting the importance of sustainable soil management practices to enhance the bioavailability of soil micronutrients [73,85,86]

### Effects of agroforestry tree species on soil quality

Results of the study showed that Calliandra and Leucaena increased soil pH above the critical level (5.5) compared to Sesbania (5.31), and indication that long term adoption of agroforestry trees has positive effects on soil quality. Similar observations were made by [66,87] who cited the effectiveness of leguminous agroforestry trees in the alleviation of soil acidity through atmospheric N_2_ fixation.

Low N contribution from agroforestry trees could be linked to the age of the tree species, low return of organic matter, and resident time of soil organic matter as affected by environmental factors such as rainfall and temperature. The enhancement of soil available P and exchangeable bases in the study under the canopy of agroforestry trees compared to soils over non-agroforestry adoption is due to the ability of trees to recycle nutrients from deeper soil layers, and the large surface area of organic matter which facilitates cation exchange and the adsorption of ions [88,89].Studies by [63,90] highlighted the ameliorative effects of Sesbania on soil available P and subsequent crop yield. Abebe et al.[4] reported the crucial role of fast-growing agroforestry shrubs and trees such as *Calliandra* and *Leucaena* in the restoration of degraded soils and replenishment of basic cations under intensive cultivation.

## 4.7 Conclusion

Findings of the study revealed that soils in Busia and Kakamega counties in western Kenya are acidic degraded as was evidenced by low concentrations of SOC (< 2.0), macronutrients (N, P, S), exchangeable bases (Ca and Mg), and micronutrients (Fe and Zn) below the critical requirements for maize production in Kenya. However, slight changes in soil parameters such as SOC content, available P, K, Mn and Cu due to agroforestry adoption highlight the potential of agroforestry adoption to enhance nutrient availability, hence soil quality. The effects of agroforestry adoption on nutrient availability and soil quality under smallholder maize production has not been fully explored due to limited use of agroforestry in soil fertility programs by small holder farmers, given competing priorities for farm resources, including fodder, biofuel, and timber at household level. It was revealed from the study that sesbania is effective in soil characteristics at both sites, whereas calliandra enhanced the physical characteristics of soils in Kakamega.

Hence, inclusion of agroforestry in extension services and soil fertility management programs is keen to conservation agriculture. However, further studies are needed to under the mechanisms of nutrient release from different agroforestry tree species and their influence on soil characteristics and maize yield under smallholder maize production in western Kenya.

## References

1. Wawire AW, Csorba Á, Tóth JA, Michéli E, Szalai M, Mutuma E, et al. Soil fertility management among smallholder farmers in Mount Kenya East region. Heliyon. 2021 Jan 2;7(3):e06488. https://linkinghub.elsevier.com/retrieve/pii/S2405844021005934

2. Abera W, Tamene L, Kassawmar T, Mulatu K, Kassa H, Verchot L, et al. Impacts of land use and land cover dynamics on ecosystem services in the Yayo coffee forest biosphere reserve, southwestern Ethiopia. Ecosyst Serv Dec 30;50:101338. https://www.sciencedirect.com/science/article/pii/S2212041621000966.

3. Verdi L, Mancini M, Ljubojevic M, Orlandini S, Dalla Marta A. Greenhouse gas and ammonia emissions from soil: The effect of organic matter and fertilisation method. Italian Journal of Agronomy. 2018 Jan 2;13(3):260–6. https://www.agronomy.it/index.php/agro/article/view/1124.

4. Abebe TG, Tamtam MR, Abebe AA, Abtemariam KA, Shigut TG, Dejen YA, et al. Growing Use and Impacts of Chemical Fertilizers and Assessing Alternative Organic Fertilizer Sources in Ethiopia. Bhatt K, editor. Appl Environ Soil Sci. 2022 Jul 3;2022:1–14. https://www.hindawi.com/journals/aess/2022/4738416/.

5. Titirmare NS, Ranshur NJ, Patil AH, Patil SR, Margal PB. Effect of Inorganic Fertilizers and Organic Manures on Physical Properties of Soil: A Review. Int J Plant Soil Sci. 2023 Jan 2;35(19):1015–23. https://journalijpss.com/index.php/IJPSS/article/view/3638

6. Autio A, Johansson T, Motaroki L, Minoia P, Pellikka P. Constraints for adopting climate-smart agricultural practices among smallholder farmers in Southeast Kenya. Agric Syst. 2021 Jul 24;194:103284. https://www.sciencedirect.com/science/article/pii/S0308521X21002377

7. Kizito F, Sène M, Dragila MI, Lufafa A, Diedhiou I, Dossa E, et al. Soil water balance of annual crop–native shrub systems in Senegal’s Peanut Basin: The missing link. Agric Water Manag. 2007 Jan 2;90(1):137–48. https://www.sciencedirect.com/science/article/pii/S0378377407000522

8. Omwakwe JA. Assessment of the Nutrient Status and Limiting Nutrients for Maize Production in Smallholder Farm Fields in Kenya. 2023 [cited 2024 Sep 6]; http://erepository.uonbi.ac.ke/handle/11295/164467

9. Nijman-Ross E, Umutesi JU, Turay J, Shamavu D, Atanga WA, Ross DL. Toward a preliminary research agenda for the circular economy adoption in Africa. Frontiers in Sustainability. 2023 Jan 2;4:1061563. https://www.frontiersin.org/articles/10.3389/frsus.2023.1061563/full

10. Kyalo D, Muyanga M, Mbuvi J, Jayne T. Geoderma The e ff ect of land use change on soil fertility parameters in densely populated areas of Kenya. Geoderma. 2019;343(January):254–62. 10.1016/j.geoderma.2019.02.033

11. Wortmann CS., Sones KR. Fertilizer use optimization in Sub-Saharan Africa. 2017;227.

12. Krah K, Michelson H, Perge E, Jindal R. Constraints to adopting soil fertility management practices in Malawi: A choice experiment approach. World Dev. 2019 Dec 30;124:104651. https://www.sciencedirect.com/science/article/pii/S0305750X19302992

13. Nyberg Y, Wetterlind J, Jonsson M, Öborn I. The role of trees and livestock in ecosystem service provision and farm priorities on smallholder farms in the Rift Valley, Kenya. Agric Syst. 2020 Jul 10;181:102815. https://linkinghub.elsevier.com/retrieve/pii/S0308521X19308789

14. Mwaura GG, Kiboi MN, Bett EK, Mugwe JN, Muriuki A, Nicolay G, et al. Adoption Intensity of Selected Organic-Based Soil Fertility Management Technologies in the Central Highlands of Kenya. Front Sustain Food Syst. 2021 Jan 2;4:570190. https://www.frontiersin.org/articles/10.3389/fsufs.2020.570190/full

15. Stewart ZP, Pierzynski GM, Middendorf BJ, Prasad PVV. Approaches to improve soil fertility in sub-Saharan Africa. Dhankher O, editor. J Exp Bot. 2020 Jan 2;71(2):632– 41. https://academic.oup.com/jxb/article/71/2/632/5581798

16. Kuyah S, Sileshi GW, Nkurunziza L, Chirinda N, Ndayisaba PC, Dimobe K, et al. Innovative agronomic practices for sustainable intensification in sub-Saharan Africa. A review. Agron Sustain Dev. 2021;41(2):1–21. http://files/1802/s13593-021-00673-4.html

17. Sileshi GW, Mpl, NAJ. Agroforestry Systems for Improving Nutrient Recycling and Soil Fertility on Degraded Lands. In: Agroforestry for Degraded Landscapes. 2020.

18. Mutuku Kinyili B. Potential of Agroforestry in Sustainable Fuelwood Supply in Kenya. Journal of Energy and Natural Resources. 2022 Jan 2;11(1):1. http://www.sciencepublishinggroup.com/journal/paperinfo?journalid=167&doi=10.11648/j.jenr.20221101.11

19. Bayala J, Kalinganire A, Sileshi GW, Tondoh JE. Soil organic carbon and nitrogen in agroforestry systems in sub-Saharan Africa: a review. Improving the profitability, sustainability and efficiency of nutrients through site specific fertilizer recommendations in West Africa agro-ecosystems. 2018;51–61. http://files/1837/978-3-319-58789-9_4.html

20. PUdawatta R, Rankoth L, Jose S. Agroforestry and Biodiversity. Sustainability. 2019 Jan 2;11(10):2879. https://www.mdpi.com/2071-1050/11/10/2879

21. Gupta SR, Dagar JC, Teketay D. Agroforestry for Rehabilitation of Degraded Landscapes: Achieving Livelihood and Environmental Security. In: Dagar JC, Gupta SR, Teketay D, editors. Agroforestry for Degraded Landscapes. Singapore: Springer Singapore; 2020. p. 23–68. http://link.springer.com/10.1007/978-981-15-4136-0_2

22. Lemage & Tsegaye. Evaluation of legume shrubs improved fallow for abandoned agricultural land rehabilitation. International Journal of Agricultural Research, Innovation and Technology (IJARIT). 2020;

23. Casanova-Lugo. Decomposition and nutrient release of leaves of tree legumes with agroforestry potential in the sub-humid tropic. Agroforest Syst. 2024;

24. Jha S, Kaechele H, Sieber S. Factors influencing the adoption of agroforestry by smallholder farmer households in Tanzania: Case studies from Morogoro and Dodoma. Land use policy. 2021 Jul 10;103:105308. https://linkinghub.elsevier.com/retrieve/pii/S0264837721000314

25. Kanyenji GM, Oluoch-Kosura W, Onyango CM, Ng’ang’a SK. Does the adoption of soil carbon enhancing practices translate to increased farm yields? A case of maize yield from Western Kenya. Heliyon. 2022 Jan 2;8(5):e09500. https://linkinghub.elsevier.com/retrieve/pii/S2405844022007885

26. Otolo JRA, Wakhungu JW. Factors influencing livelihood zonation in Kenya. International Journal of Education and Research. 2013 Jul 10;1(12):1–10. http://www.ijern.com/journal/December-2013/24.pdf

27. Nyberg Y, Jonsson M, Laszlo Ambjörnsson E, Wetterlind J, Öborn I. Smallholders’ awareness of adaptation and coping measures to deal with rainfall variability in Western Kenya. Agroecology and Sustainable Food Systems. 2020 Jul 10;44(10):1280–308. https://www.tandfonline.com/doi/full/10.1080/21683565.2020.1782305

28. Nachtergaele FO. Classification Systems: FAO ?. In: Reference Module in Earth Systems and Environmental Sciences. Elsevier; 2017. p. B9780124095489106000. https://linkinghub.elsevier.com/retrieve/pii/B9780124095489105202

29. Omuto CT. Major Soil and Data Types in Kenya. In: Developments in Earth Surface Processes. Elsevier; 2013. p. 123–32. https://linkinghub.elsevier.com/retrieve/pii/B9780444595591000116

30. Cheboiwo JK, Langat D, Mutiso F, Cherono F. A Review Farm Forestry Evolution for the Last 100 Years in Kenya: A Look at Some Key Phases and Driving Factors. 2024 Jul 11; https://core.ac.uk/reader/234656560

31. Okalebo JR, Gathua KW, Woomer PL. Laboratory methods of soil and plant analysis: a working manual second edition. Sacred Africa, Nairobi. 2002 Jul 11;21:25–6.

32. Bouyoucos GJ. Hydrometer Method Improved for Making Particle Size Analyses of Soils1 | Agronomy Journal. 2024. https://acsess.onlinelibrary.wiley.com/doi/10.2134/agronj1962.00021962005400050028x

33. Robinson JBD. Tropical Soil Biology and Fertility: A Handbook of Methods (Second Edition). Edited By J. M. Anderson and J. S. I. Ingram, with 13 appendices by various authors. Wallingford, Oxfordshire: CAB International (1993), pp. 221, £19.95. ISBN 0-85198-821-0. Exp Agric. 1994 Jul 12;30(4):487. https://www.cambridge.org/core/product/identifier/S0014479700024832/type/journal_article

34. Hazelton & Murphy. Interpreting Soil Test Results: What Do All the Numbers Mean? CSIRO Publishing, Australia. 2007;CSIRO Publishing, Australia, 1–152.

35. NAAIAP & KARI. Soil Suitability Evaluation for Maize Production in Kenya. Nairobi; 2014.

36. Nelson DW and LES. Total Carbon, Organic Carbon, and Organic Matte. In: Methods of soil analysis. Madison: ASA and SSSA; 1996. p. 961–1010.

37. E.O. Mclean. Soil pH and Lime Requirement. In: Methods of Soil Analysis: Part 2 Chemical and Microbiological Properties, 922, Second Edition. Agronomy Monographs; 1982. p. 200–10.

38. J. M. Bremner. Nitrogen-Total. Soil Science Society of America. 1996;1085–120.

39. Olsen SR and DLA. Phosphorus, Methods of Soil Analysis Part 2, Chemical and Microbiological Properties. Madison, Wisconsin: American Society of Agronomy, Inc. 1965;

40. Sumner ME, & MWP. Cation exchange capacity and exchange coefficients. 1996.

41. Fox R. L., Evaluating the Sulfur Status of Soils by Plant and Soil Tests. Soil Science of America Journal. 1964;28(2).

42. Norvell WA. Norvell, W.A. (1984). Comparison of chelating agents as extractants for metals in diverse soil materials. Soil Science Society of America Journal, 1984;48(6):1285–1292.

43. Syano NM, Moses M. Nyangito MN, Kironchi G, Wasonga OV. Agroforestry practices impacts on soil properties in the drylands of Eastern Kenya. Trees, Forests and People. 2023;

44. Kinyili B, Ndunda E, Kitur E. Influence of Duration of Agroforestry on Physico-Chemical Soil Quality Parameters in Machakos County, Kenya. 2021;

45. Abera. Soil bulk density, soil moisture content and yield of Tef (Eragrostis tef) influenced by Acacia seyal Del canopy in Parkland agro-forestry system. J Soil Sci Environ Manage. 2019;10(6):124–9.

46. Biswas S, Hazra GC. Establishment of critical limits of indicators and indices of soil quality in rice-rice cropping systems under different soil orders. Geoderma. 2017;34– 48.

47. Silva G. Soil physical quality of Luvisols under agroforestry, natural vegetation and conventional crop management systems in the Brazilian semi-arid region. Geoderma. 2011;61–70.

48. Hairiah K. Soil carbon stocks in Indonesian (agro) forest transitions: Compaction conceals lower carbon concentrations in standard accounting. Agric Ecosyst Environ. 2020;

49. Throop HL. When bulk density methods matter: Implications for estimating soil organic carbon pools in rocky soils. 2012;66–71.

50. Udawatta R, Anderson SH. CT-measured pore characteristics of surface and subsurface soils influenced by agroforestry and grass buffers. Geoderma. 2008;381–9.

51. P. Udawatta R Rljs. Agroforestry and Biodiversity. Sustainability. 2019;

52. Gama-Rodrigues EF. Carbon Storage in Soil Size Fractions Under Two Cacao Agroforestry Systems in Bahia, Brazil. Environmental Management. Brazil Environmental Management. 2010;45(2):274–83.

53. Hairiah K, van Noordwijk M, Sari RR, Saputra DD, Widianto Suprayogo D, et al. Soil carbon stocks in Indonesian (agro) forest transitions: Compaction conceals lower carbon concentrations in standard accounting. Agric Ecosyst Environ. 2020 Jun 1;294.

54. Udawatta RP, Anderson SH, Gantzer CJ, Garrett HE. Agroforestry and Grass Buffer Influence on Macropore Characteristics. Soil Science Society of America Journal. 2006 Sep;70(5):1763–73.

55. Kinyili Ndunda, E.N., & Kitur, E. (2020). BM. Influence of Duration of Agroforestry on Physico-Chemical Soil Quality Parameters in Machakos County, Kenya. Journal of Applied Science, Engineering and Technology for Development. 2020 Aug 12; https://jasetd.dkut.ac.ke/publication/view

56. Nyuma HT, Churu H. A review on challenges and opportunities in management of soils of arid and semi-arid regions of Kenya. East African Journal of Environment and Natural Resources. 2022 Jul 3;5(1):303–17. https://journals.eanso.org/index.php/eajenr/article/view/840

57. Zuza, E, Maseyk K, Bhagwat S, Factors affecting soil quality among smallholder macadamia farms in Malawi. Agriculture & Food Security volume. 2023;12(7).

58. Nguru WM, Gachene CK, Onyango CM, Ng’ang’a SK, Girvetz EH. Factors constraining the adoption of soil organic carbon enhancing technologies among small-scale farmers in Ethiopia. Heliyon. 2021 Nov 27;7(12).

59. Maier R, Schack-Kirchner H, Nyoka IB, Lang F. Gliricidia intercropping supports soil organic matter stabilization at Makoka Research Station, Malawi. Geoderma Regional. 2023;35.

60. Suleiman A, Yit AT, Pfeifer M, Galloway J, Otene I. Effects of Tree Planting on Soil Organic Carbon and Nitrogen Content in Nigerian Agroforestry Systems. Medicon Agriculture & Environmental Sciences. 2023;

61. Pankaj, Bhardwaj KK, Yadav R G. Role of agroforestry systems in enrichment of soil organic carbon and nutrients: A review. ECJ. 2023;25(1):289–96.

62. Hijbeek R, et al. Liming agricultural soils in Western Kenya: Can long-term economic and environmental benefits pay off short term investments? Agric Syst. 2021;190:1030–5.

63. Rautaray SK, Pradhan S, Mohanty S et al. Energy efficiency, productivity and profitability of rice farming using Sesbania as green manure-cum-cover crop. Nutr Cycl Agroecosyst. 2020;83–101.

64. Simelane MP, S., Soundy P,, Maboko MM. Effects of Rainfall Intensity and Slope on Infiltration Rate, Soil Losses, Runoff and Nitrogen Leaching from Different Nitrogen Sources with a Rainfall Simulator. Sustainability. 2023;

65. Sahu H, Ku, MS, MAP, & KS. Impact of organic and inorganic farming on soil quality and crop productivity for agricultural fields: A comparative assessment. Environmental Challenges. 2024;

66. Rizwan. Changes in soil chemical properties in agroforestry system at USU Arboretum Kwala Bekala. Earth Environ Sci. 2020;

67. Fahad S. Agroforestry Systems for Soil Health Improvement and Maintenance. Sustainability. 2022;

68. Riyadh Z, Rahman M, Saha S, Ahamed T CD. Adaptation of agroforestry as a climate smart agriculture technology in Bangladesh. Int J Agril Res Innov & Tech. 2021;49– 59.

69. Fahad S, Chavan SB, Chichaghare AR, Uthappa AR, Kumar M, Kakade V, et al. Agroforestry Systems for Soil Health Improvement and Maintenance. Sustainability 2022, Vol 14, Page 14877. 2022 Nov 10 [cited 2024 Sep 6];14(22):14877. https://www.mdpi.com/2071-1050/14/22/14877/htm

70. Muchane MN, Sileshi GW, Gripenberg S, Jonsson M, Pumariño L, Barrios E. Agroforestry boosts soil health in the humid and sub-humid tropics: A meta-analysis. Agric Ecosyst Environ. 2020 Jun 15;295.

71. Mabagala FS, Mng’ong’o ME. On the tropical soils; The influence of organic matter (OM) on phosphate bioavailability. Vol. 29, Saudi Journal of Biological Sciences. Elsevier B.V.; 2022. p. 3635–41.

72. Neina D, Adolph B. Sulphur Contents in Arable Soils from Four Agro-Ecological Zones of Ghana. Land (Basel). 2022 Oct 1;11(10).

73. Njoroge R, Otinga AN, Okalebo JR, Pepela M, Merckx R. Maize (Zea mays L.) Response to secondary and micronutrients for profitable n, p and k fertilizer use in poorly responsive soils. Agronomy. 2018;8(4).

74. Paramisparam P, Ahmed OH, Omar L, Ch’ng HY, Johan PD, Hamidi NH. Co-application of charcoal and wood ash to improve potassium availability in tropical mineral acid soils. Vol. 11, Agronomy. MDPI; 2021.

75. Diallo MB, Akponikpè PBI, Fatondji D, Abasse T, Agbossou EK. Long-term differential effects of tree species on soil nutrients and fertility improvement in agroforestry parklands of the Sahelian Niger. Forests Trees and Livelihoods. 2019;28(4):240–52. 10.1080/14728028.2019.1643792

76. Kachaka ÉY, Munson AD, Gélinas N, Khasa D. Adoption of an improved fallow practice using Acacia auriculiformis on the Batéké Plateau in the Democratic Republic of the Congo. Agroforestry Systems. 2020 Jun 1;94(3):1047–58.

77. Shin R. Strategies for Improving Potassium Use Efficiency in Plants. Mol Cells. 2014;575–84.

78. Kaur M, Singh S, Dishri M, Singh G, Singh SK. Foliar application of zinc and manganese and their effect on yield and quality characters of potato (Solanum tuberosum L.) cv. Kufri Pukhraj. Plant Arch. 2018;18(2):1628–30.

79. Michael PS. Role of Organic Fertilizers in the Management of Nutrient Deficiency, Acidity, and Toxicity in Acid Soils-a Review. Journal of Global Agriculture and Ecology. 2021;12(2):19–30.

80. Abindaw T, Hanyabui E, Atiah K, Akwasi EA, Ziblim IA. Influence of land use types on the distribution of selected soil properties in tropical soils of the Coastal Savanna zone. Heliyon. 2023 Mar 1;9(3).

81. Yadav MBN, Patil PL, Hebbara M. Comparative Studies on Soil Quality Index Estimation of a Hilly-Zone Sub-Watershed in Karnataka. Sustainability. 2023;

82. Noulas C, Tziouvalekas M, Karyotis T. Zinc in soils, water and food crops. Journal of Trace Elements in Medicine and Biology. 2018;252–60.

83. Van Eynde E, Breure MS, Chikowo R et al. Soil zinc fertilisation does not increase maize yields in 17 out of 19 sites in Sub-Saharan Africa but improves nutritional maize quality in most sites. Plant Soil. 2023;67–91.

84. Hernandez-Soriano, Jimenez-Lopez. Linking dissolved organic matter composition to metal bioavailability in agricultural soils: effect of anionic surfactants. Biogeosciences Discuss. 2015;12:5697–723.

85. Kihara J, Manda J, Kimaro A, Swai E. Contributions of integrated soil fertility management (ISFM) to various sustainable intensification impact domains in Tanzania. Agric Syst. 2022;203.

86. Ngaba MJY, Mgelwa AS, Gurmesa GA, Uwiragiye Y, Zhu F, Qiu Q, et al. Meta-analysis unveils differential effects of agroforestry on soil properties in different zonobiomes. Plant Soil. 2023 Jan 3; https://link.springer.com/10.1007/s11104-023-06385-w

87. Krah K, Michelson H, Perge E, Jindal R. Constraints to adopting soil fertility management practices in Malawi: A choice experiment approach. World Dev. 2019 Dec 1;124.

88. Zhang Y, Liu X, Zhang C, Lu X. A combined first principles and classical molecular dynamics study of clay-soil organic matters (SOMs) interactions. Geochim Cosmochim Acta. 2020;291:110–25.

89. Strawn DG. Sorption Mechanisms of Chemicals in Soils. Soil Syst. 2021;5(13).

90. Palsaniya DR, Dhyani SK, Rai P. Silvipasture in India present perspectives and challenges ahead. Vol. 137. Scientific Publishers; 2011. 207 p. https://www.scientificpub.com/upload/pdf/143.pdf

